# Mutations in *GPNMB* associated with Amyloid cutis dyschromica alter intracellular trafficking and processing of GPNMB

**DOI:** 10.1101/2023.12.19.572409

**Authors:** Erin C. Bogacki, Patrick A. Lewis, Susanne Herbst

## Abstract

Amyloid cutis dyschromica (ACD) is a rare skin condition characterized by focal areas of hyperpigmentation with hypopigmented macules and distinct regions of amyloid deposition. Until recently, the genetic cause of ACD remained unknown. Several studies have since named GPNMB truncation mutations as causal, with protein loss-of-function underlying ACD pathogenesis. GPNMB missense mutations have also been observed in patients, but these are less well characterized; especially on a cellular level. Here, we observed that GPNMB missense mutations implicated in familial ACD show distinct cellular phenotypes that result in impaired protein maturation and processing from the endoplasmic reticulum (ER) to the *trans*-Golgi network (TGN), prompting failed trafficking to lysosomes. These missense mutations also show failed secretion of the extracellular fragment of GPNMB, a well-characterized property of the protein. Overall, this work highlights previously undescribed cellular characteristics of GPNMB missense mutations implicated in ACD and helps to better inform the clinically observed phenotypes, as well as underscore GPNMB’s role at melanosomes.

## 1. Introduction

Amyloidosis cutis dyschromica (ACD), first described by Morishima in 1970 (Kuseyri et al. 2017), is a skin condition characterized by patterned spots of hyperpigmentation with hypopigmented macules and focal areas of keratinocyte-derived amyloid deposition (Mahon et al. 2016; Yang et al. 2018). Primarily affecting East and Southeast Asian individuals, ACD is a rarely documented variant of primary localized cutaneous amyloidosis (PLCA) that manifests in individuals during adolescence. There is typically a large gap between the age of onset and time of diagnosis, as symptoms are often unremarkable except for mild puritis in approximately 19 % of cases (Mahon et al. 2016). Until recently, its genetic cause remained unknown; however, Mahon et al. (Mahon et al. 2016) emphasized genetic involvement was very likely, as most cases were familial, and equally affected both males and females. While sporadic cases do exist, they are rare, and age of onset is largely seen into adulthood. The genetic cause of ACD has only been recently elucidated. Yang et al. (Yang et al. 2018) conducted whole-exome sequencing of an affected family and identified two candidate genes, NIPAL2 on chromosome 8 and GPNMB on chromosome 7, as potentially causative. Across all nine studied individuals, six mutations predicted to cause truncation of GPNMB were identified, of which existed in extremely low frequencies in ethnicity-matched controls. Therefore, premature termination of the GPNMB protein caused by mutations in the GPNMB gene was named as causative of ACD, with complex modes of inheritance (i.e., both autosomal dominant and recessive inheritance, and semi-dominant inheritance in heterozygous mutation carriers) (Yang et al. 2018; Qin et al. 2021). Since its identified role in ACD, several missense point mutations have been found: p.G363V and p.I174M point mutations in a cohort study of an affected British family with Pakistani ancestry \cite{Rahman2021-ur} and an autosomal dominant p.C413S point mutation has been described in individuals of Filipino and Han Chinese origin (Onoufriadis et al. 2019; Qin et al. 2021).

Glycoprotein nonmetastatic protein B (GPNMB) is a type I transmembrane glycoprotein first isolated from metastatic melanoma cells (Weterman et al. 1995). Encoded by the GPNMB gene at locus 7p15, it has assumed several names in accordance with the tissue type it has been isolated from (i.e., osteoactivin in osteoclasts, hematopoietic growth factor inducible neurokinin-1 in kidneys)(Safadi et al. 2001; Bandari et al. 2003). It consists of several domains: the extracellular domain contains an RGD integrin-binding motif, a polycystic kidney disease (PKD) domain, and a KLD domain that mediates tumorigenicity (Tomihari et al. 2009; Theos et al. 2013; Xie et al. 2019; Tsou and Sawalha 2020; Han et al. 2021). The intracellular domain contains various sorting motifs, including a lysosomal sorting motif (Hoashi et al. 2010). GPNMB is a heavily glycosylated protein that contains 12 putative glycosylation sites (Tsou and Sawalha 2020; Han et al. 2021). Glycosylation in the endoplasmic reticulum (ER) results in a GPNMB precursor protein that is further glycosylated within the trans-Golgi network (TGN) resulting in its mature form (Hoashi et al. 2010). Additionally, the extracellular domain can be cleaved and secreted into the extracellular environment (Rose et al. 2010). The function of GPNMB remains elusive. It is widely expressed across various tissues (e.g., skin, brain, muscle, and bone) and is under the transcriptional regulation of the TFEB family of transcription factors, implying a function for GPNMB in the endolysosomal system (Gutknecht et al. 2015; Biswas et al. 2020; Carey et al. 2020; Ripoll et al. 2008). Indeed, GPNMB has been linked to the lysosome upon induction of lysosomal stress. Specifically, previous work has shown that GPNMB is preferentially expressed in response to various instances of lysosomal dysfunction, such as pH neutralization and lipid overload (Gabriel et al. 2014; Moloney et al. 2018; Van Der Lienden et al. 2018). In the context of ACD, skin pigmentation requires the formation of melanosomes, a type of lysosome related organelles (LROs), in melanocytes. Because melanosomes are integral to the maintenance of melanin deposition throughout the skin, mutations in genes that encode melanosome-associated proteins can result in altered pigment deposition (Le et al. 2021). GPNMB is highly expressed in melanocytes and localizes to melanosomes but is also expressed to a lesser degree on the surface of cells; promoting adhesion to keratinocytes via its RGD motif (Tomihari et al. 2009). GPNMB has been found on melanosomes at all stages, indicating a potential role in their development (Chi et al. 2006; Zhang et al. 2012). Knockdown of GPNMB has shown to reduce melanosome formation in all stages, further highlighting its role in the formation of early melanosomes and pigmentation (Zhang et al. 2012). Therefore, the phenotype observed in ACD cases, if GPNMB is vital to melanosome function and pigmentation, is not unexpected.

GPNMB truncation mutations indicate that GPNMB loss-of-function underlies ACD pathogenesis, however the impact of GPNMB point mutations on GPNMB function and ACD pathogenesis is less clear. This study aimed to characterize the described missense point mutations for potential GPNMB loss of function. We show that ACD-related point mutations result in abnormal ER and Golgi processing of GPNMB, which results in GPNMB mislocalization and impaired lysosomal recruitment. Additionally, we demonstrate that ACD-related point mutations result in impaired secretion of the GPNMB extracellular domain. Taken together, these findings suggest that the so far reported ACD-related GPNMB point mutations result in a loss of GPNMB function in accordance with reported truncation mutants, further emphasizing GPNMB function in melansosome health and the etiopathogenesis of ACD.

## 2. Materials and Methods

### 2.1 Plasmids

The open reading frame of full-length human GPNMB (NM_001005340.2) cloned into pcDNA3.1-C-EGFP was purchased from GenScript. pcDNA3.1-GPNMB-EGFP-G375V, GPNMB-EGFP-I174M, and GPNMB-EGFP-C425S constructs were generated by site-directed mutagenesis using a Q5 Site-Directed Mutagenesis Kit (#E0552S, New England Biolabs). All plasmids were sequenced to verify successful modification. DNA constructs were replicated in *E. coli* DH5α (#11583117, Fisher Scientific) and extracted using a QIAprep Spin Miniprep Kit (Qiagen).

### 2.2 Antibodies

Anti-GPNMB (AF2550) from R&D Systems, anti-β-Actin (clone AC-15) from Sigma-Aldrich, and anti-GFP (#MA5-15256) from ThermoFisher Scientific were used for Western blotting. Anti-LAMP1 (#H4A3) from Developmental Studies Hybridoma Bank, and anti-TGN46 (#13573-1-AP) from Proteintech were used for immunofluorescence. Secondary antibodies for Western Blotting were anti-goat and anti-mouse-HRP coupled (Sigma-Aldrich) and for immunofluorescence were anti-rabbit-AF568 and anti-mouse-AF647 (ThermoFisher Scientific).

### 2.3 Cell culture

293T endothelial-like cells (ATCC CRL-3216) were obtained from ATCC. HEK 293T cells were grown in DMEM with added 10% heat-inactivated FCS and maintained in an incubator at 37 degrees Celsius and 5% CO_2_. Cells were seeded at a concentration of 3 x 10^5^ cells/well in 12-well plates for Western blotting, 1.25 x 10^5^ cells/well on coverslips coated in poly-d-lysine (Gibco, ThermoFisher Scientific) in 24-well plates for immunofluorescence, and 4.0 x 10^4^ cells/well for ELISA protocols.

### 2.4 Plasmid transfection

HEK293T cells were seeded on 24- or 12-well plates and incubated overnight. The cells were transfected with either wildtype GPNMB or G375V, I174M, or C425S plasmid constructs using FuGENE HD Transfection Reagent (#E2311, Promega) at a DNA:FuGENE ratio of 1:3. Cells were transfected with 0.5 µg or 1 µg plasmid DNA for 24-well or 12-well plates respectively. For 96-well plates, cells were reverse-transfected with 125 ng of plasmid DNA.

### 2.5 LLOMe treatment

L-leucyl-L-leucine methyl ester (LLOMe; BAChem, #4000725.0001) was stored at a stock concentration of 333 mM in ethanol and diluted to 1 mM in 10% FCS in DMEM for experimental use. Cells were administered LLOMe for 1 hour, after which it was removed and cells were processed for immunofluorescence.

### 2.6 Western blotting

Cells were lysed on ice in 1 % Triton-X-100/Tris lysis buffer (9803S, Cell Signaling) containing a protease and phosphatase inhibitor cocktail (78442, Thermo Scientific). Samples were spun for 15 minutes at 16,200 g at 4 °C. Post-nuclear supernatant was collected and stored at -20 ° C. Samples were denatured at 80 °C for 8 minutes in LDS sample buffer containing 1x NuPAGE reducing agent (ThermoFisher Scientific). Samples were run on a NuPAGE 4-12 % Bis-Tris gel (ThermoFisher Scientific) and transferred to a PVDF membrane using a Turbo-Blot transfer system (Biorad). The PVDF membrane was blocked for 1 hour in a TBS-T-milk solution (5 % skimmed milk powder, TBS, 0.05 % Tween-20). After blocking, membranes were incubated at 4 °C overnight with primary antibodies diluted 1:1000 in TBS-T-milk solution, and then secondary antibodies diluted 1:10’000 in TBS-T-milk solution for 45 minutes at room temperature. Western blots were developed using an iBright imaging system (ThermoFisher Scientific) and quantified with densitometry using FIJI software (Schindelin et al. 2012). A detailed protocol can be found under http://dx.doi.org/10.17504/protocols.io.4r3l22yz4l1y/v1.

### 2.7 Immunofluorescence

Cells seeded in 24-well plates on poly-D-Lysine-coated glass coverslips were washed once in PBS and fixed with 4 % methanol-free PFA (Electron Microscopy Science) for 15 minutes at 4 °C. Coverslips were blocked and permeabilized in 0.3 % Triton-X-100/5 % FCS/PBS solution for 20 minutes at room temperature. Coverslips were incubated for 1 hour at room temperature with primary antibodies diluted 1:100 in 5 % FCS/PBS. Samples were then washed 3 times with PBS followed by incubation with secondary antibodies diluted 1:1000 in 5 % FCS/PBS. Coverslips were washed 3 times with PBS again and nuclei were stained in 300 nM DAPI for 10 minutes. Samples were washed a final time and mounted to microscope slides using DAKO mounting medium. Images were taken using a Leica SP8 inverted confocal microscope. A detailed protocol can be found under http://dx.doi.org/10.17504/protocols.io.4r3l22yz4l1y/v1.

### 2.8 ELISA

Supernatant was collected 48 hrs post transfection and 18 hours after the last medium change. All supernatant was stored at -80 °C until ready for processing. Live cells were counterstained with Hoechst and EGFP fluorescence was imaged on a Spark Cyto plate reader (Tecan) to estimate transfection efficiency. GPNMB levels in the supernatant were measured by ELISA (DuoSet ELISA Development system, Human Osteoactivin/GPNMB, #DY2550, R&D Systems) according to the manufacturer’s instructions.

## 3. Results

### 3.1 C425S and I174M *GPNMB* missense mutations result in abnormal protein processing and N-glycosylation

*GPNMB* missense mutations are found throughout the extracellular domain of *GPNMB* with I174M localizing to the N-terminal domain, G375V to the PKD domain and C425S to the KLD domain (Figure 1A). Mapping these mutations on the predicted GPNMB Alphafold (Jumper et al. 2021) structure indicated that I174M might disturb an intra-molecular interaction in the N-terminal domain. G375V is found in a region of the protein for which no high-confidence folding prediction is available, therefore hindering a prediction of the impact of the mutation on protein function. In contrast, C425 constitutes one of six highly-conserved Cysteine residues commonly found in Kringle domains. These cysteines form three disulfide bonds which are crucial for the correct folding of the domain (Chrystal et al. 2021) (Figure 1B).

**Figure 1.**
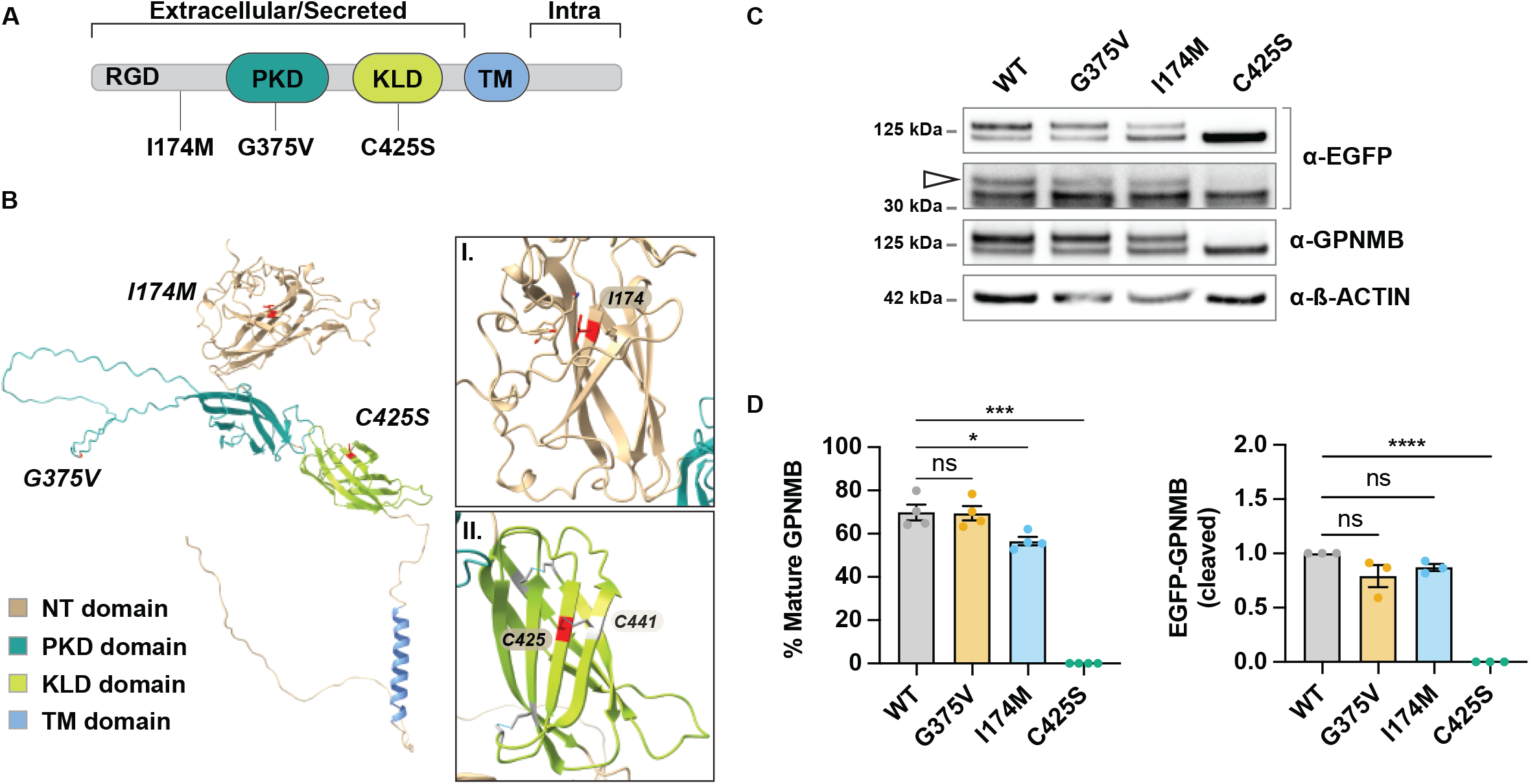
Location of GPNMB ACD mutations and their effects on GPNMB protein expression and maturation. **A**. Cartoon figure illustrating GPNMB domain structure and the location of associated ACD mutations. RGD = Arginyl-Glycyl-Aspartic acid motif, PKD = Polycystic Kidney Disease domain, KLD = Kringle-like domain, TM = Transmembrane domain. **B**. Alphafold predicted structure of GPNMB featuring NT, PKD, KLD, and TM domains. Missense mutations implicated in GPNMB are labeled within their respective regions. Insets I. and II. show interactions of I174 and C425. **C**. HEK293 cells transfected with wildtype GPNMB-EGFP or G375V, I174M, and C425S mutant GPNMB-EGFP were analyzed for GPNMB mature (M) and precursor protein (P1) levels. A representative Western blot is shown. The arrow indicates the cleaved GPNMB C-terminal fragment. **D**. Data show the percentage of the mature form of GPNMB out of total GPNMB protein (M + P1) and the quantification for the cleaved C-terminal GPNMB fragment. Data represent the mean ± SEM of three to four independent biological replicates. *p < 0.05, ***p<0.001, ****p < 0.0001, ns: not significant by ordinary one-way ANOVA.

Previous work has shown that GPNMB is post-translationally modified through N-glycosylation of its ectodomain, occurring in both the ER (precursor (P1) form) and trans-Golgi network (mature (M) form)(Hoashi et al. 2010). To determine the effects of ACD missense mutations on the maturation of the protein, HEK cells were transfected with either wildtype pcDNA3.1-GPNMB-EGFP or mutant constructs corresponding to ACD mutations: pcDNA3.1-GPNMB-EGFP-G375V, GPNMB-EGFP-I174M, and GPNMB-EGFP-C425S and differences in GPNMB protein levels, specifically of its precursor and mature forms, were analyzed by Western blot using an antibody which targets the N-terminus of GPNMB. We observed that, in cells expressing wildtype GPNMB, the M form of the protein made up approximately 74 % of total observed GPNMB (M + P1; Figure 1C-D). There was no significant difference observed in the G375V mutant compared to wildtype. The I174M mutant resulted in a significant (∼13.5 %) reduction of the M form of GPNMB, indicating a failure of trans-Golgi processing. The C425S mutant resulted in a complete loss of the M form, indicative of total processing failure from the ER to the Golgi. We also assessed GPNMB cleavage by monitoring the presence of the cleaved C-terminal low-molecular-weight fragment by using an anti-GFP antibody which recognizes the C-terminal EGFP-tag of the GPNMB construct used. We did not observe any major differences in GPNMB cleavage except for the C425S mutant which showed a complete loss of the cleavage product (Figure Figure 1C-D).

### *GPNMB* missense mutations implicated in ACD result in impaired trafficking and lysosomal targeting of the protein

Because of GPNMB’s association with endolysosomal function, we sought to investigate the cellular localization of the protein upon induction of lysosomal damage, and whether this would be altered in ACD mutants. Wildtype GPNMB and G375V, I174M, and C425S mutant GPNMB-expressing HEK cells were treated with either control medium or the lysosomal damaging agent LLOMe for 1 hour. At steady-state levels, GPNMB was found primarily at the TGN. Upon stimulation with LLOMe, wildtype GPNMB localised to LAMP-1 positive structures, indicating lysosomal recruitment in response to lysosomal stress (Figure 2). Lysosomal localization of GPNMB was, however, impaired in the ACD mutants. In GPNMB-I174M transfected cells, some colocalization with LAMP-1 was observed, but to a lesser degree than wildtype. In G375V mutant cells, there was no significant increase in GPNMB localization to lysosomes in LLOMe conditions compared to control. Additionally, GPNMB colocalized with TGN46 signal, a trans-Golgi marker, indicating impaired maturation and trafficking from the trans-Golgi network to lysosomes in both G375V and I174M mutants. C425S mutant GPNMB showed complete failure of lysosomal targeting, and instead presented a reticular phenotype; indicative of defective trafficking from the ER.

**Figure 2.**
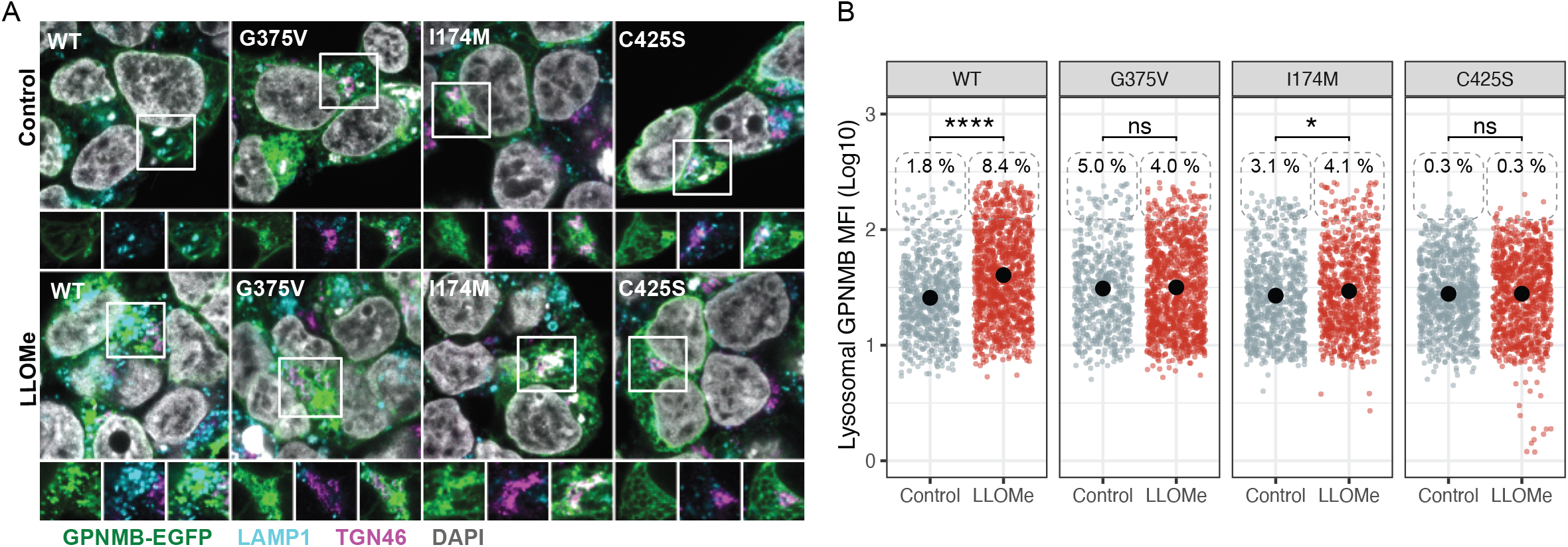
Intracellular localization of GPNMB missense mutants. HEK293 cells transfected with GPNMB-EGFP constructs were stimulated with LLOMe (1 mM) for 1 hr. Lysosomes were labeled with LAMP-1 and the TGN with TGN46. **A**. Representative images of GPNMB localization. **B**. Quantification of the lysosomal GPNMB mean fluorescence intensity (MFI) indicative of GPNMB recruitment and calculation of % of GPNMB positive lysosome. The black dot corresponds to the mean. *p < 0.05, **** p < 0.0001, ns: not significant by Wilcoxon rank sum test.

### 3.3 Secretion of the GPNMB extracellular domain is reduced in ACD-associated *GPNMB* missense mutant cells

GPNMB ectodomain shedding in human melanocytes was first observed *by Hoashi et al*. (Hoashi et al. 2010). Similarly, Rose *et al*. (Rose et al. 2010) determined that cleavage of the extracellular domain of GPNMB in breast cancer cells is mediated by the ADAM10 protease. We therefore aimed to determine if this property of GPNMB is altered in cells expressing ACD-associated mutant GPNMB. HEK cells expressing wildtype GPNMB and either G375V, I174M, or C425S mutant GPNMB were cultured over-night and secretion of GPNMB into the supernatant was measured by ELISA. Cells expressing mutant GPNMB, specifically the G375V and C425S mutants, showed significantly reduced GPNMB secretion. In contrast, secretion of the GPNMB I174M mutants was comparable or higher than wildtype (Figure 3A). All GPNMB constructs were expressed to comparable levels (Figure 3B).

**Figure 3.**
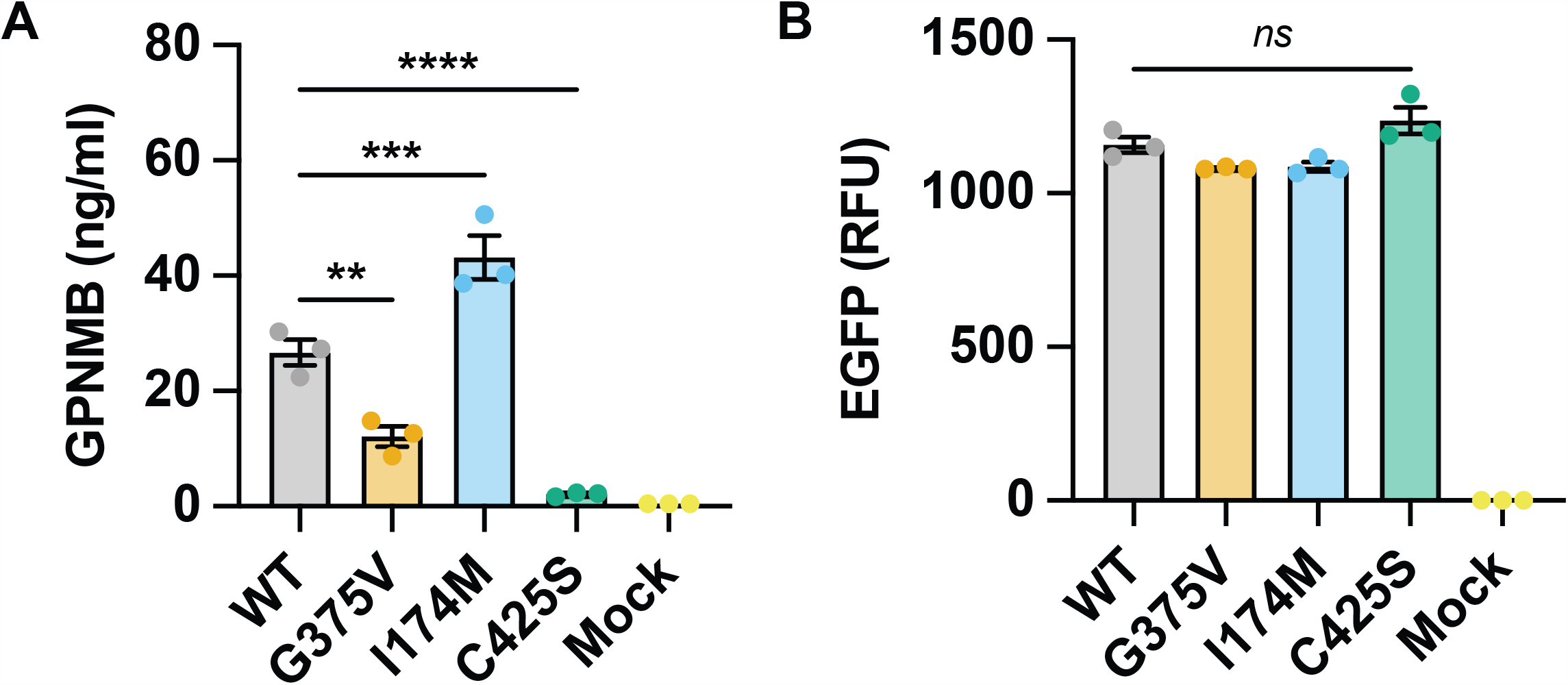
Effect of missense mutations on GPNMB secretion. HEK293 cells transfected with GPNMB-eGFP missense mutants were analyzed for GPNMB secretion by ELISA. **A**. Data show the quantification of GPNMB secreted into supernatant and represents one experiment out of 3 biological replicates. **B**. Quantification of GPNMB-EGFP transfection efficiency was estimated as EGFP relative fluorescence units (RFU) per cell. Mock denotes non-transfected HEK293 cells. **p < 0.01, ***p < 0.001, ****p < 0.0001, ns: not significant by ordinary one-way ANOVA.

Taken together, these results indicate that reported GPNMB ACD missense mutations result in impaired GPNMB maturation and secretion, indicative of loss-of-function of the protein.

## 4. Discussion

While a causal role for GPNMB mutations in ACD has been recently elucidated, little has been shown regarding how point mutations may alter GPNMB function on a cellular level. Here, we show that GPNMB point mutations implicated in familial ACD result in impaired trafficking and processing of the protein. Previous work has highlighted an important function of GPNMB at melanosomes, and our data suggests a molecular phenotype of GPNMB missense mutants that could underlie the clinical phenotypes observed in ACD patients.

Our data suggests that the C425S GPNMB mutant is only present in the P1 form, which precedes the mature form of the protein. This indicates that either Golgi N-glycosylation of GPNMB does not occur, or that this mutant form of GPNMB is not shuttled beyond the ER for further downstream processing and secretion. Exit of proteins from the ER is dependent upon proper protein folding, and those that are incorrectly folded are often retained within the organelle (Kincaid and Cooper 2007). According to the Alphafold predicted structure of GPNMB, the C425 cysteine forms a Cys-Cys bond with the cysteine located at C457 (Figure 1B). The C425S mutation implicated in ACD would, therefore, disrupt this bond; leading to misfolding of GPNMB. This is further corroborated by the intracellular localisation of the C425S mutant, which showed a reticular pattern that did not overlap with either the TGN and lysosomes, and the complete lack of secretion. Together, these findings indicate that the C425S mutant is retained in the ER and constitutes a loss-of-function variant.

The G375V and I174M GPNMB mutants, while implicated in ACD, showed less severe cellular phenotypes than the C425S mutant. While there was some overlap in GPNMB-EGFP signal with TGN46, indicative of GPNMB that remains within the trans-Golgi network, there was still observable colocalization of I174M mutant with LAMP-1; therefore, lysosomal localization of GPNMB still occurs. Some failure of GPNMB trafficking from the TGN is evident in both mutants, but these point mutations do not seem to fully impair GPNMB function in our model, as both formed punctate structures that localized to lysosomes to a degree. Similarly, the G375V mutant showed ratios of P1 and M forms of protein comparable to wildtype GPNMB. These differences in GPNMB molecular phenotype may be due to the way they affect protein processing and folding, but this requires further investigation.

The shedding and secretion of GPNMB’s ectodomain is increasingly becoming a property of interest for several fields of research. We, therefore, determined if the G375V, I174M, or C425S mutants implicated in ACD showed changes in GPNMB secretion. While the I174M mutant showed secretion comparable to wildtype GPNMB, both the G375V and C425S mutants showed reduced levels of GPNMB secretion. Cleavage and secretion of the extracellular fragment occurs at the cell membrane, where the mature form of the protein is trafficked after secretion from the trans-Golgi network. If, as our data suggests, the mutants fail to surpass the ER and Golgi, whether partially or completely, it is not unexpected that GPNMB secretion fails.

Here, we have observed distinct cellular phenotypes of GPNMB missense mutations implicated in ACD, suggesting varying levels of severity dependent upon the point mutation. As there is currently no data to suggest that ACD patients with these mutations present with different clinical phenotypes, it would be of value to determine if these mutations do result in varying degrees of clinical severity of ACD. Overall, this work suggests that GPNMB missense mutations implicated in ACD result in loss-of-function phenotypes similar to those seen in GPNMB truncation mutations, and provides insights into the cellular dysfunction that underlies clinical observations of ACD. We suggest that ACD GPNMB point mutations result in failure to properly process and traffic GPNMB, resulting in impaired melanosome maturation.

## Abbreviations

(ACD): Amyloid cutis dyschromica
(PLCA): Primary localized cutaneous amyloidosis
(GPNMB): Glycoprotein nonmetastatic protein B
(NIPAL2): NIPA like domain containing 2
(RGD): Arginyl-glycyl-aspartic acid
(PKD): Polycystic kidney disease
(KLD): Kringle-like domain
(TFEB): Transcription factor EB
(LROs): Lysosome related organelles
(MITF): Microphthalmia-associated transcription factor
(TM): Transmembrane
(ADAM10): A disintegrin and metalloproteinase domain-containing protein 10
(Di-L): Di-leucine
(EGFP): Enhanced green fluorescent protein
(FCS): Fetal calf serum
(PBS): Phosphate-buffered saline
(LLOMe): L-leucyl-L-leucine methyl ester
(PFA): Paraformaldehyde
(DAPI): 4′,6-diamidino-2-phenylindole
(BSA): Bovine serum albumin
(ER): Endoplasmic reticulum
(TGN): trans-Golgi network

## 5. Acknowledgements

This research was funded by Aligning Science Across Parkinson’s (ASAP 000478) through the Michael J. Fox Foundation for Parkinson’s Research (MJFF). For the purpose of open access, the authors have applied a CC BY public copyright license to all Author Accepted Manuscripts arising from this submission.

## 6. Data and materials availability

Raw data presented in this study have been deposited in Zenodo (doi 10.5281/zenodo.10255251) and all plasmids generated in this study are available upon request from the corresponding authors.

## RESOURCES TABLE

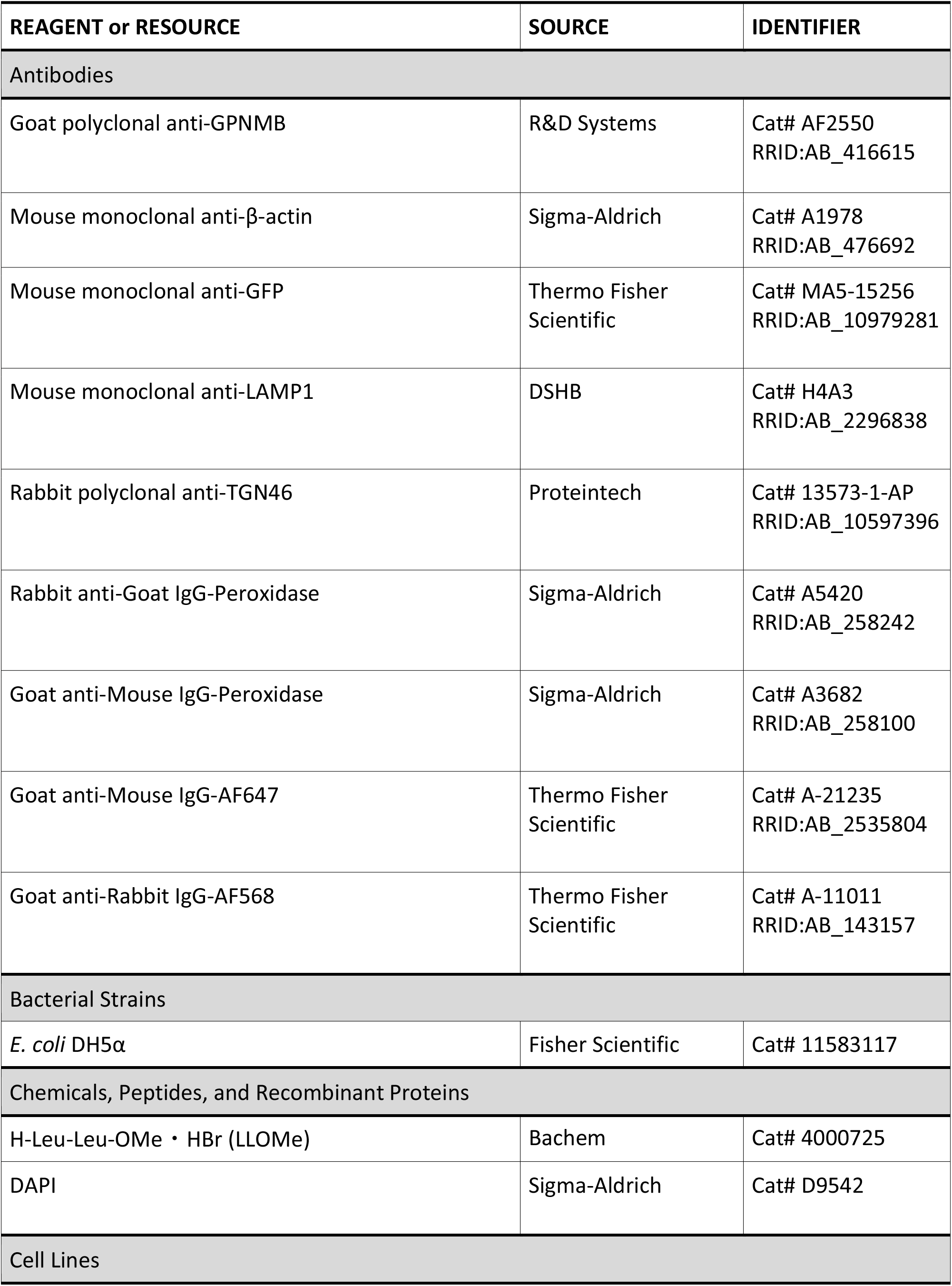
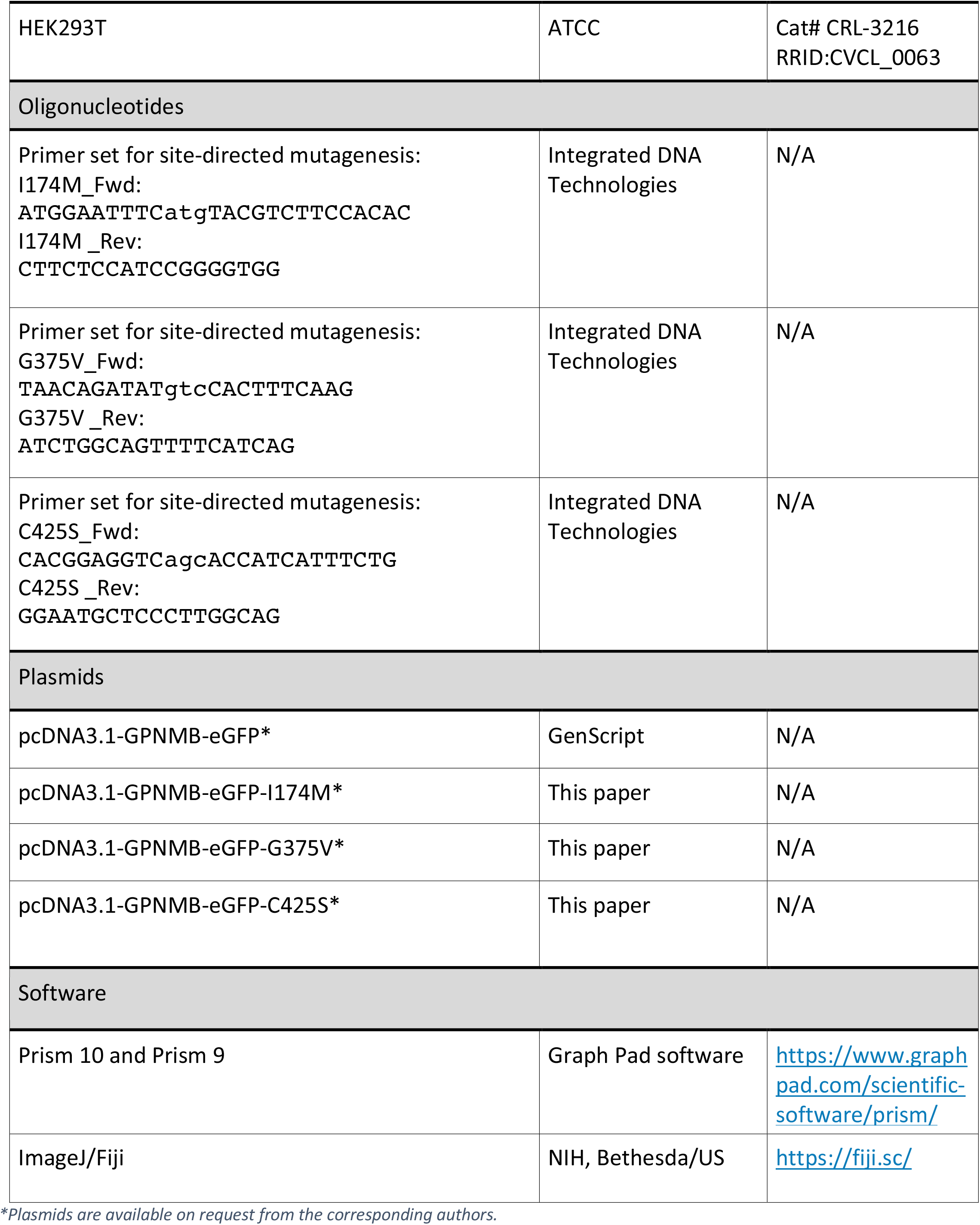

## Notes

### Competing Interest Statement

The authors have declared no competing interest.

